# Emergent dynamics of underlying regulatory network links EMT and androgen receptor-dependent resistance in prostate cancer

**DOI:** 10.1101/2022.12.06.516625

**Authors:** Rashi Jindal, Abheepsa Nanda, Maalavika Pillai, Kathryn E Ware, Divyoj Singh, Manas Sehgal, Andrew J. Armstrong, Jason A Somarelli, Mohit Kumar Jolly

## Abstract

Advanced prostate cancer patients initially respond to hormone therapy, be it in the form of androgen deprivation therapy or second-generation hormone therapies, such as abiraterone acetate or enzalutamide. However, most men with prostate cancer eventually develop hormone therapy resistance. This resistance emerges in several ways, such as through genetic mutations, epigenetic mechanisms, or through non-genetic pathways, such as lineage plasticity along epithelial-mesenchymal or neuroendocrine-like axes. These mechanisms of hormone therapy resistance often co-exist within a single patient’s tumor and can overlap within a single cell. There exists a growing need to better understand how phenotypic heterogeneity and plasticity results from emergent dynamics of the regulatory networks governing androgen independence. Here, we investigated the dynamics of a regulatory network connecting the drivers of androgen receptor (AR) splice variant-mediated androgen independence and those of epithelial-mesenchymal transition. Model simulations for this network revealed four possible phenotypes: epithelial-sensitive (ES), epithelial-resistant (ER), mesenchymal-resistant (MR) and mesenchymal-sensitive (MS), with the latter phenotype occurring rarely. We observed that well-coordinated “teams” of regulators working antagonistically within the network enable these phenotypes. These model predictions are supported by multiple transcriptomic datasets both at single-cell and bulk levels, including *in vitro* EMT induction models and clinical samples. Further, our simulations reveal spontaneous stochastic switching between the ES and MR states. Addition of the immune checkpoint molecule, PD-L1, to the network was able to capture the interactions between AR, PD-L1, and the mesenchymal marker SNAIL, which was also confirmed through quantitative experiments. This systems-level understanding of the driver of androgen independence and EMT could aid in understanding non-genetic transitions and progression of such cancers and help in identifying novel therapeutic strategies or targets.

## Introduction

Hormone therapy resistance in prostate cancer often arises as a consequence of androgen-independent activation of pro-survival androgen receptor (AR) signaling (1). Several genetic alterations are associated with AR re-activation during ADT, such as amplification of AR, AR structural rearrangements (2), and gain-of-function mutations in the AR ligand binding domain (3). In addition to genetic mechanisms of AR-reactivation, hormone therapy-resistant prostate cancer also exhibits an increase in specific splice variants of full-length AR (AR-FL) (4). The canonical AR gene has 8 exons that code for four different domains: the N-Terminal Domain (NTD), DNA-binding domain (DBD), hinge region, and Ligand Binding Domain (LBD). AR-V7 is a splice variant that lacks the LBD but can translocate into the nucleus (5). AR-V7 has been implicated in hormone therapy resistance in preclinical models (6,7,8,9) and has been shown to be predictive of shorter time to progression and poorer overall survival in men with advanced prostate cancer (10,11).

In addition to these AR-dependent mechanisms, AR-independent mechanisms of resistance also exist, such as bypass signalling via the glucocorticoid receptor signalling pathway (12) and the emergence of aggressive variant prostate cancer, mediated through lineage plasticity toward AR-low or AR-null, neuroendocrine-like (11,13), FGFR-driven (14), and/or epithelial-mesenchymal transition (EMT)-like phenotypes (15,16).

EMT, a cell biological process implicated in wound healing and embryogenesis, is often aberrantly activated in cancer, and can promote migration, invasion, and survival during metastatic dissemination. Once the disseminating cancer cells reach a distant organ through the bloodstream, they usually undergo the reverse of EMT – Mesenchymal-Epithelial Transition (MET) – to facilitate colonization (17). EMT also promotes immune suppressive and immune-evasive phenotypes through both secretion of extracellular immune-suppressive cytokines, such as IL-6, IL-8, and GROα (reviewed in (18)), and by direct transcriptional activation of immune checkpoints, such as PD-L1 (19,20). The EMT-induced alterations in immune signalling and function may also induce a positive feedback toward a more EMT-like state (reviewed in (18)). In addition to its role in metastasis and immune suppression/evasion, EMT and therapeutic resistance have been found to be linked across multiple drugs and cancer types (21,22). Despite the relationships between EMT-like biology and hormone therapy resistance, the crosstalk between EMT and AR signaling remains poorly understood. For instance, while EMT is enriched in hormone therapy-resistant prostate cancer, overexpression of AR, especially that of AR-V7, has also been shown to increase levels of genes associated with stemness and EMT (23). Conversely, the EMT factor, Snail, induces resistance to enzalutamide through alterations in AR signaling (15,24), suggesting that in prostate cancer, the EMT and AR-FL/AR-V7 axes can influence one another. AR-FL and AR-V7 can upregulate mesenchymal markers, such as SNAI1, ZEB1, fibronectin, N-cadherin and vimentin (23,24). Further, knockdown of AR in C4-2 cells disrupted the ability of drugs such as enzalutamide to induce EMT. This effect was mediated by the direct transcriptional control of EMT-inducing transcription factor SNAI1 (Snail) by AR (25), implying that modulation of AR activity can, in specific lineage contexts, direct EMT-like lineage plasticity. Moreover, in bone metastases from castrationresistant prostate cancer (CRPC), immunohistochemical analysis revealed upregulation of EMT-related proteins, suggesting potential clinical implication of EMT in CRPC (26). In human C4-2B CRPC xenograft models knockdown of ISL1, a driver of EMT, attenuated enzalutamide resistance and reduced tumor growth (27), thereby highlighting putative bidirectional interplay between EMT and AR signaling pathway (28). EMT also promotes immune suppressive and immune-evasive phenotypes through both secretion of extracellular immune-suppressive cytokines, such as IL-6, IL-8, and GROα(18), and by direct transcriptional activation of immune checkpoints, such as PD-L1 (19,20). The EMT-induced alterations in immune signalling and function may also induce a positive feedback toward a more EMT-like state (18). This interplay between multiple signalling axes can lead to an aggressive phenotype with enhanced migration, invasion, immune evasive capacity, and increased survival during metastatic dissemination.

Here, we elucidate the emergent dynamics of the interconnection between the EMT and AR axes through mathematical modeling of experimentally defined and curated underlying regulatory networks. Dynamical simulations for these networks revealed two dominant phenotypes: epithelial-sensitive (ES) and mesenchymal-resistant (MR), indicating that EMT and CRPC may promote each other. These model-based predictions were validated by analysis of multiple publicly available preclinical and clinical datasets at bulk and singe-cell level. Finally, we integrated PD-L1 in our network to demonstrate how the three axes of immune-evasion, androgen-independence and EMT may be interconnected. These results offer mechanistic insights into empirical observations about EMT and CRPC and demonstrate how non-genetic/non-mutational changes, such as phenotypic plasticity, are capable of aggravating disease progression.

## Results

### Crosstalk between EMT and AR signaling pathways can lead to multiple phenotypes

We identified a regulatory network that encompasses the experimentally-reported interactions among various key components of EMT (miR-200, ZEB1, SLUG, SNAIL) and AR signaling (AR, AR-V7) pathways as well as factors known to connect these axes (hNRNPA1, LIN28, let-7; **Fig 1A, i; Table S1**). The connections included in this network were obtained from multiple *in vitro* and *in vivo* experimental data sets, such as knockdown or overexpression, ChIP-seq, and protein-RNA interaction data. For instance, miR-200 and ZEB1 can inhibit each other (29,30), as can LIN28 and let-7 (31), ZEB1 and AR (32), and SNAIL and SLUG (33). SNAIL and SLUG can both activate AR-V7 (15,34), which can activate LIN28 (23). These interactions lead to the formation of various interconnected feedback loops, capable of enabling complex emergent dynamics of this regulatory network. We have used such experimentally-informed approach earlier to uncover novel insights about the dynamics of EMT and its association with drug resistance in ER+ breast cancer (33,35).

**Figure 1.**
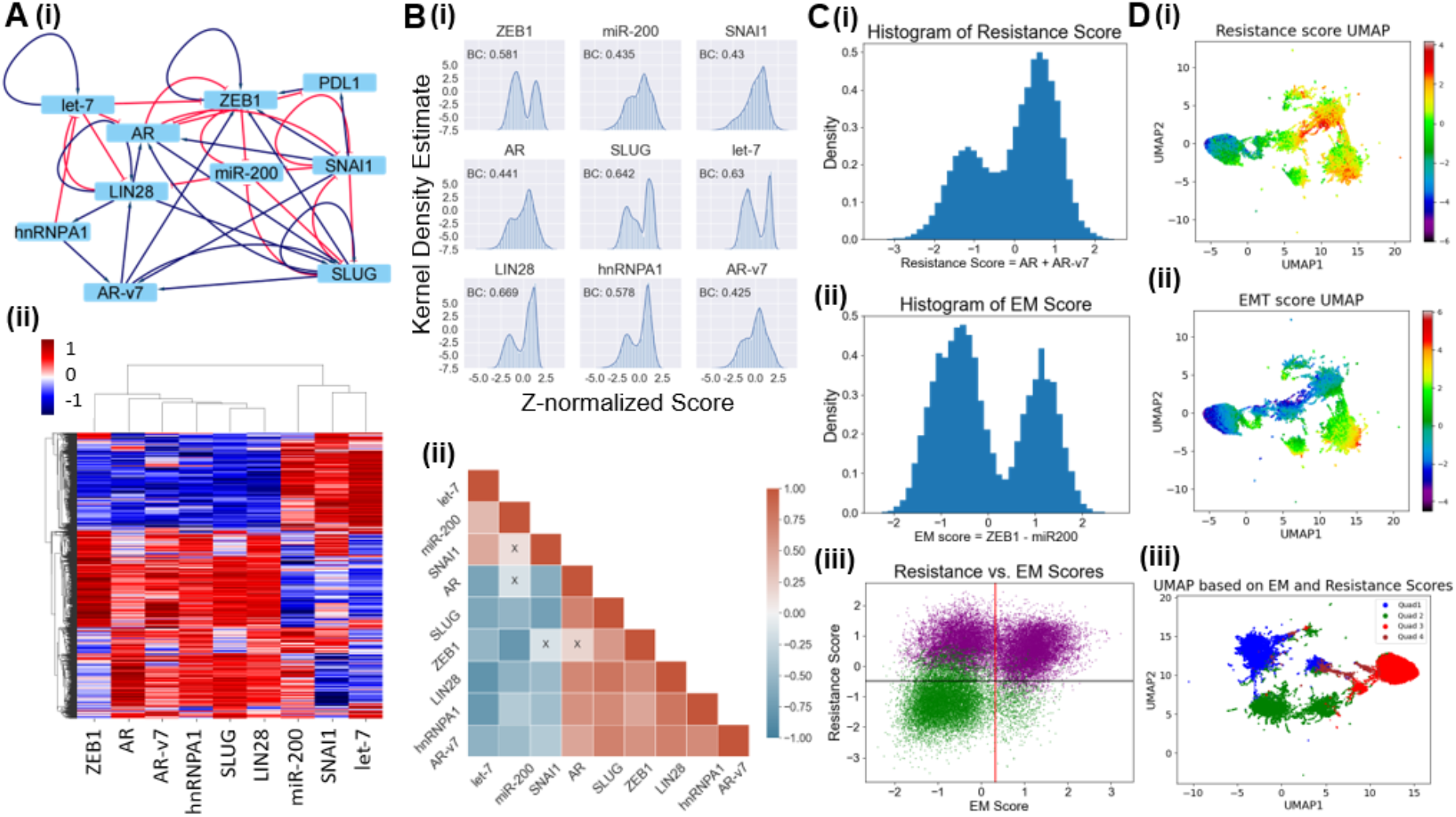
Multi-stable dynamics of the coupled EMT – AR crosstalk network. **A) (i)** Gene regulatory network (GRN) showing crosstalk between EMT and Androgen Receptor (AR, AR-v7) signalling. Blue arrows represent activation links; red hammers represent inhibition. **(ii)** Heatmap of stable steady-state solutions for network shown in A (i), as obtained via RACIPE. **B) (i)** Kernel Density Estimate plots with histograms of z-score levels of individual nodes in a network, fitted to Gaussian distributions. Each panel legend shows the corresponding Bimodality Coefficient (BC). **(ii)** Pairwise correlation matrix in which each cell denotes the correlation coefficient between the corresponding set of genes. Red indicates positive correlation; blue indicates negative correlation, Colourmap indicates the strength and sign of correlation; cells with a cross (x) highlight nonsignificant correlation (p >= 0.05). **C) (i)** Histogram of Resistance score (AR+AR-V7) fit to a Gaussian. **(ii)** Histogram of epithelial-mesenchymal (EM) score (ZEB1 - miR200) fit to a Gaussian. **(iii)** Scatter plot showing the corresponding EM and resistance scores for all RACIPE solutions obtained. UMAP (uniform manifold approximation and projection) dimensionality reduction plots for steady-state solutions states colored by Resistance score; colormap indicates the Resistance score. **(ii)** Same as (i) but for EM score. (iii) UMAP based on EM and resistance scores together; Quadrant 1 represents Mesenchymal Resistant, Quadrant 2 represents Epithelial Resistant, Quadrant 3 represents Epithelial Sensitive and Quadrant 4 represents Mesenchymal Sensitive populations.

To decode the emergent dynamics of this network, the network was simulated using RACIPE (Random Circuit Perturbation) (36) which converts a network topology into a set of coupled ordinary differential equations (ODEs), and simulates the dynamics for an ensemble of kinetic parameters chosen from a biologically-relevant range. Thus, each unique parameter set in the context of RACIPE can be thought of as representing an individual cell in a population, displaying cell-to-cell variability. Such variability can lead to fractional killing and determine the IC50 of a population to a given drug (37). For each parameter set, it further samples various initial conditions to identify different possible steady states (phenotypes) onto which the system can converge.

For the given gene regulatory network, we obtained multiple stable states that can be visualized as a hierarchically-clustered heatmap (**Fig 1A, ii**). From a qualitative perspective, miR-200, SNAIL and let-7 are co-expressed in this heatmap; similarly, ZEB1, SLUG, LIN28, AR, AR-V7 and hNRNPA1 are often co-expressed. To quantify these co-expression patterns, we calculated the bimodality coefficients for distributions of steady state values for each of these network components. The kernel density estimates (KDE) plots with histogram fits show that the mesenchymal markers, such as ZEB1 and SLUG, stemness factors, such as LIN28 and let-7, and hnRNPA1 show clearly bimodal patterns; these trends were not as clearly observed for levels of miR-200, SNAIL, AR and AR-V7 (**Fig 1B,i; Table S2, S3).** Next, to investigate the co-expression patterns in the heatmap more quantitatively, we calculated the pairwise correlation coefficients among all pairs of network nodes, across all steady state values given by RACIPE (**Fig 1B, ii**). In this pairwise correlation matrix, we observed two groups – or “teams” – of factors, such that members in one “team” were all positively correlated with one another, but negatively correlated with members of the other “team”(38), thus reminiscent of patterns seen in earlier analysis of networks underlying phenotypic plasticity in cancer (20,39,40). Here, one “team” is comprised of let-7, miR-200 and SNAIL while the other team contains AR-V7, SLUG, LIN28 and hnRNPA1 as its members. These two “teams” can act antagonistically, forming a “toggle switch”, thus driving phenotypic heterogeneity, which is observed in the case of EMT-CRPC crosstalk.

We next sought to define the phenotypes along the epithelial-mesenchymal and drug resistance axes. To do this, we defined an ‘EM score’ and an AR-mediated ‘Resistance score’ for our simulation results. Higher EM scores (difference between the z-normalized expression of ZEB1 and miR-200) indicate more mesenchymal samples. Resistance score was defined as the sum of z-normalized expression of AR and AR-V7, based on evidence that shows that higher expression of both AR and its splice variant AR-V7 can mediate the emergence of AR-dependent mechanisms of hormone therapy resistance in multiple experimental settings (41,42). The Resistance score in this context does not consider AR-independent models of hormone therapy resistance, such as neuroendocrine-like phenotypes(43) or “double negative” (AR-/NEPC-) phenotypes (14). A higher Resistance score indicates a stronger AR-mediated therapy resistance. The histograms for both these scores revealed bimodality (**Fig 1C, i–ii**). Next, to observe these phenotypic combinations, we plotted a scatter plot with the EM scores and Resistance scores as axes **(Fig 1C,iii)**. Points are coloured according to K-means clustering, with the highest silhouette score obtained for K=2 **(Fig S1A)**. After identifying cut-offs for EM and Resistance scores as obtained from respective bimodal histograms, three key phenotypes are observed: epithelial/ sensitive (ES), mesenchymal/resistant (MR) and epithelial/resistant (ER). The scatter plot reveals that K-means algorithm clusters the ER and MR phenotypes together, indicating that although some cells may not necessarily undergo a canonical EMT, they may still show signs of hormone therapy resistance. The mesenchymal/sensitive (MS) phenotype was also apparent at a minor frequency, indicating that most of the prostate cancer cells undergoing EMT are likely to show phenotypic traits consistent with hormone therapy resistance.

The points in each quadrant were then mapped onto a UMAP plot, with colours of points representing different phenotypes **(Fig 1D)**. Here again we observe that the different phenotypes form different clusters, with MS less well-delineated as a unique cluster, likely due to its low occurrence. Together, investigation of these dynamical properties of the EM and Resistance axes suggests that the dominant phenotypes for this network are ER, MR and ES.

### Emergence of co-existing phenotypes is driven by coordinated “teams” within a network

The heatmap and correlation matrix obtained after RACIPE simulation highlight the presence of two groups of co-expressed genes (**Fig 1A,ii; 1B,ii**): one of them comprising ZEB1, SLUG, AR, AR-V7 and hnRNPA1, and the other comprising let-7, miR-200 and SNAIL. The first group contains EMT-inducing transcription factors in PCa (ZEB1, SLUG) (44,45), as well as drivers of ADT resistance (AR, AR-V7 and hnRNPA1) (46). Except for SNAIL, the second group contains EMT-inhibiting microRNAs (let-7, miR-200). In our model, we have considered one node to represent multiple members of the miR-200 family, most members of which can inhibit and are inhibited by ZEB1 and ZEB2 (47) and can inhibit EMT and metastasis in prostate cancer (48,49). Together, these two groups may be reflective of the two major phenotypes illuminated by our simulations: 1) the group with upregulated ZEB1, SLUG, AR, AR-V7 and SLUG corresponding to a MR phenotype, and 2) the group with downregulated levels of these molecules reflecting an ES phenotype. It is worth noting that these models likely do not include scenarios of lineage plasticity toward AR null lineage states and are focused on models that maintain AR expression.

Next, we asked whether emergence of these two groups of co-expressed molecules could be an outcome of specific kinetic parameter values chosen and sampled by RACIPE, or whether they are attributed to a consequence of the underlying topology of the regulatory network irrespective of parameters. To interrogate this further, we mapped the “influence matrix” corresponding to the simulated network. This matrix is defined purely on the basis of network topology without simulating network dynamics, as previously defined (39). Each element in this matrix defines how strongly a node in the network (represented by cell in the i^th^ row) influences the levels of another node in the network (represented by cell in the j^th^ column), through multiple paths of varying lengths through which these two nodes are connected in the given network. The influence matrix corresponding to our biological network **(Fig 2A)** reveals that the network is comprised of two “teams”, such that members across teams are mutually inhibitory, but those within a team activate each other. The composition of these two “teams” is identical to that of two groups of co-expressed molecules (**Fig 1B, ii**): one “team” including ZEB1, SLUG, AR, AR-V7 and SLUG, and the other including AR, AR-V7 and hnRNPA1. This analysis suggests that the underlying network topology defines the two “teams”.

**Figure 2.**
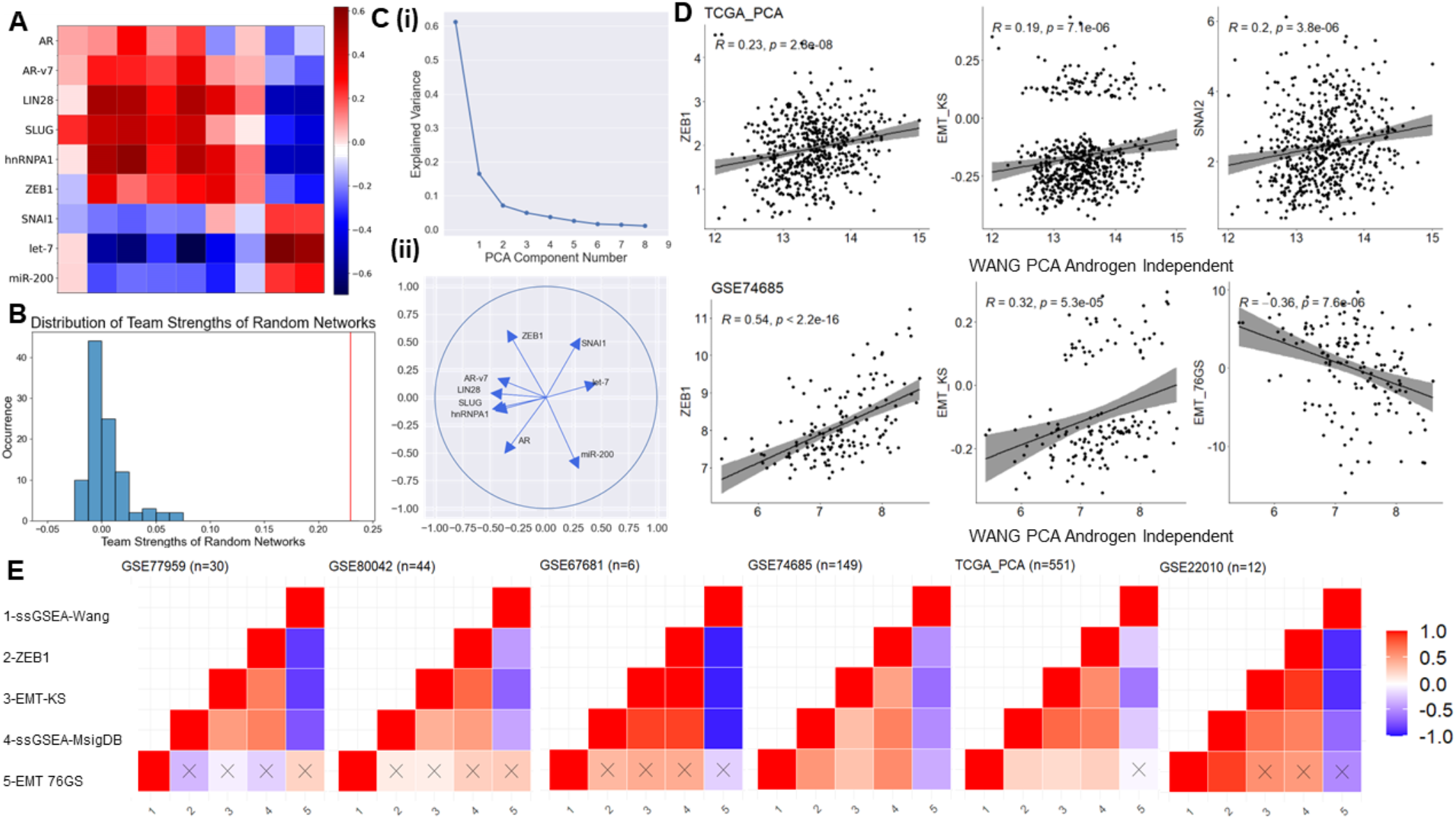
Coordinated EMT and AR programs in network topology and transcriptomic data. **A)** Influence matrix for the biological network shown in Fig 1A. **B)** Histogram of group strengths for 100 random (non-biological) networks generated by shuffling edges in the biological network. The group strength of the biological network us shown in red **C) (i)** Scree plot showing PCA components and their explained variance percentage **(ii)** PCA Correlation Circle of the RACIPE solutions of the biological network **D)** Scatter plots indicating Spearman’s correlation between metrics of EMT (EMT KS score, EMT 76GS score, ZEB1 and SNAI2 expression levels) and ssGSEA scores for Wang Prostate Cancer Androgen Independent gene set, in TCGA (above) and GSE74685 (below). **E)** Pairwise correlation between different EMT scoring and androgen independence gene lists/ metrics in different transcriptomic datasets: (Left to right - GSE77959 (n=30), GSE80042 (n=44), GSE67681 (n=6), TCGA (n=551) and GSE22010 (n=12)). Labels: 1 – ssGSEA scores for Wang Prostate Cancer Androgen Independent geneset (MsigDB), 2 – ZEB1 expression levels, 3 – KS score, 4 – ssGSEA scores for Hallmark EMT geneset (MsigDB), 5 – 76GS score. Colour bar indicates Spearman’s correlation coefficient. Crosses indicate corresponding p-value >= 0.05.

We next interrogated whether the presence of “teams” is specific to the network considered here. To address this question, we calculated the “team strength” of the “teams” identified in the influence matrix. This metric quantifies the strength of the two teams in the influence matrix by providing an estimate of how strongly members within a team support each other and how strongly members across teams inhibit each other. This estimate is defined on a scale of [0, 1], with a higher corresponding to more well-defined teams. We first generated an ensemble of 100 randomized (hypothetical) networks with the same number of nodes and edges, but varied network topology **(Fig S1D)** and calculated their “team strength”. While the calculated “team strength” of the randomly generated “teams” was largely centered around 0, the team strength of the experimentally-derived network was 0.2297 **(Fig 2B)**, indicating that the presence of “teams” in the experimentally-informed network is largely unique to the topology of underlying network. Moreover, in the ensemble of random networks generated, the higher the team strength, the stronger the correlation between EMT and Resistance scores, suggesting that the presence of these “teams” contributes directly to the functional coupling between these two axes of plasticity (**Fig S1B–C**).

To confirm the composition of the “teams” obtained via an influence matrix, we performed principal component analysis (PCA) on the ensemble of solutions obtained by RACIPE simulations. The Scree plot **(Fig. 2C, i)** shows that 61.21% of the variance can be explained by PC1 and 16.45% of it can be explained by PC2, thus first 2 PCs can explain about 80% variance. To better visualize this trend, we created a correlation circle **(Fig 2C, ii)** that shows that members from one “team” were located close to each other on one side of the circle, while those from the other team located close to each other on the other side of it. This observation further consolidates the proposed mutually antagonistic relationship between these two “teams” of molecules embedded in underlying network topology. The mutual antagonism between these “teams” control the various phenotypes observed in prostate cancer along the EMT and therapeutic resistance axes, for instance, as showcased by the presence of miR200 in one team and ZEB1 in the opposite team. A comparative analysis of this wild-type (WT) network with its randomized counterparts revealed that the formation of “teams”, as identified by PCA plot, is specific to WT network (**Fig S2**). Overall, this analysis reveals that “teams” containing interconnected molecular players underlie observed association between EMT and androgen independence.

### Clinical data supports model predictions about the association of EMT with therapy resistance

To provide additional support for the association between EMT and AR signaling – as indicated by the predominance of ES and MR phenotypes – we examined how EMT and AR signatures correlated in various clinical data sets. One of the datasets (GSE74685) contained transcriptomic data from 62 patients with castration-resistant prostate cancer (26), while another cohort (n = 149) was obtained from The Cancer Genome Atlas (TGCA). We quantified the single-sample Gene Set Enrichment Analysis (ssGSEA) scores based on the Wang Androgen Independent Prostate Cancer (29) geneset from Molecular Signature Database (MSigDB) (50). In both GSE74685 and TCGA cohorts, these ssGSEA scores were found to correlate positively with expression levels of ZEB1 and SNAI2, both of which promote a mesenchymal phenotype. Besides individual molecules, EMT scores based on larger gene sets – KS and 76GS (51) – showed similar trends. The higher the KS score or the lower the 76GS score, the more mesenchymal a sample is. Consistent with model predictions, the ssGSEA score for the androgen independence gene list correlated positively with KS and negatively with 76GS **(Fig 2D)**.

To further validate our model predictions, we analyzed the following additional transcriptomic datasets – GSE77959 (human prostate specimens) (52), GSE80042 (time-course *in vitro* EMT induction in LNCaP cells) (45), GSE67681 (cells from mice with *Pten* deletion and *Kras* activation) (53), GSE74685 (17) (CRPC tumors) and GSE22010 (immortalized prostate epithelial cells) (55). Analysis of pairwise correlations among the ssGSEA scores for the androgen independence gene set, ZEB1 expression levels, KS and 76GS scores and ssGSEA scores for the Hallmark EMT gene set revealed positive correlations between the ssGSEA score for androgen independence and ZEB1, KS and the ssGSEA score for Hallmark EMT, and all four of these metrics correlated negatively with 76GS EMT scores **(Fig 2E)**. Together, these results suggest that AR-dependent models of androgen independence and EMT status are tightly coupled.

### Stochastic stimulations demonstrate cell-state switching among interconnected EMT and resistance traits

Stochasticity in various biological processes, such as transcription and translation, can produce non-genetic heterogeneity (56,57). These stochastic factors can drive phenotypic switching, especially in a multi-stable system (58–60), such as the coupled EMT-AR circuit. Thus, it is possible that a cell can reversibly switch to a drug-tolerant state in response to drug treatment, and later lead to repopulation and consequent tumor relapse. Hence, we investigated the effect of noise/stochasticity on the dynamics of our gene regulatory network. We identified parameter sets via RACIPE that resulted in multi-stable states and then performed stochastic simulations using specific parameter sets.

Among the ensemble of parameter sets in RACIPE simulations, 21.3% were monostable, 27.8% were bistable, 25.8% were tri-stable and 25.1% had more than three states. Hence, the network considered here is predominantly multi-stable; in approximately 80% of cases, the system could converge to more than one stable state (phenotype), depending on the initial simulations. Among the different possible combinations of states observed in bistable parameter sets, the most common combination (37.21%) was the co-existence of ES (epithelial-sensitive) and MR (mesenchymal-resistant) (**Fig 3A**). From the bistable parameter sets corresponding to ES and MR phenotypes, we simulated the system in the absence of noise for multiple initial conditions and plotted the EM Score and Sensitivity scores. Sensitivity score here is defined as the negative of the resistance score, i.e., a higher Sensitivity score means a Sensitive phenotype while a lower Sensitivity score means a Resistance phenotype. These simulations showed two distinct levels of steady-state EM scores and Sensitivity scores (**Fig 3B),** each corresponding to the Epithelial and Mesenchymal region for the EM plot and Resistant and Sensitive for the Sensitivity plot. We also observed that the initial conditions that lead to an Epithelial state on the EM plot also led to Sensitive state on the Sensitivity plot. The same was also true for the mesenchymal and resistant state, thus supporting the associated dynamics and crosstalk of EMT and AR signaling. Next, we simulated the system dynamics under the influence of noise in gene expression. We observed that the system switched from mesenchymal-resistant (MR) to epithelial-sensitive (ES) phenotype in the presence of gene expression noise. This switch along the EM scores and sensitivity scores happened almost simultaneously, thereby indicating a strong coupling of EMT and hormone therapy resistance axes in our GRN (**Fig 3C**). To strengthen our analysis, we simulated the system under the presence of gene expression noise for multiple initial conditions and plotted the landscape of observed trajectories. We observed that the system revealed two deep “attractors” corresponding to ES and MR states (**Fig 3D**), thus enabling co-existence of these two cell-states in a population of heterogeneous gene expression.

**Figure 3:**
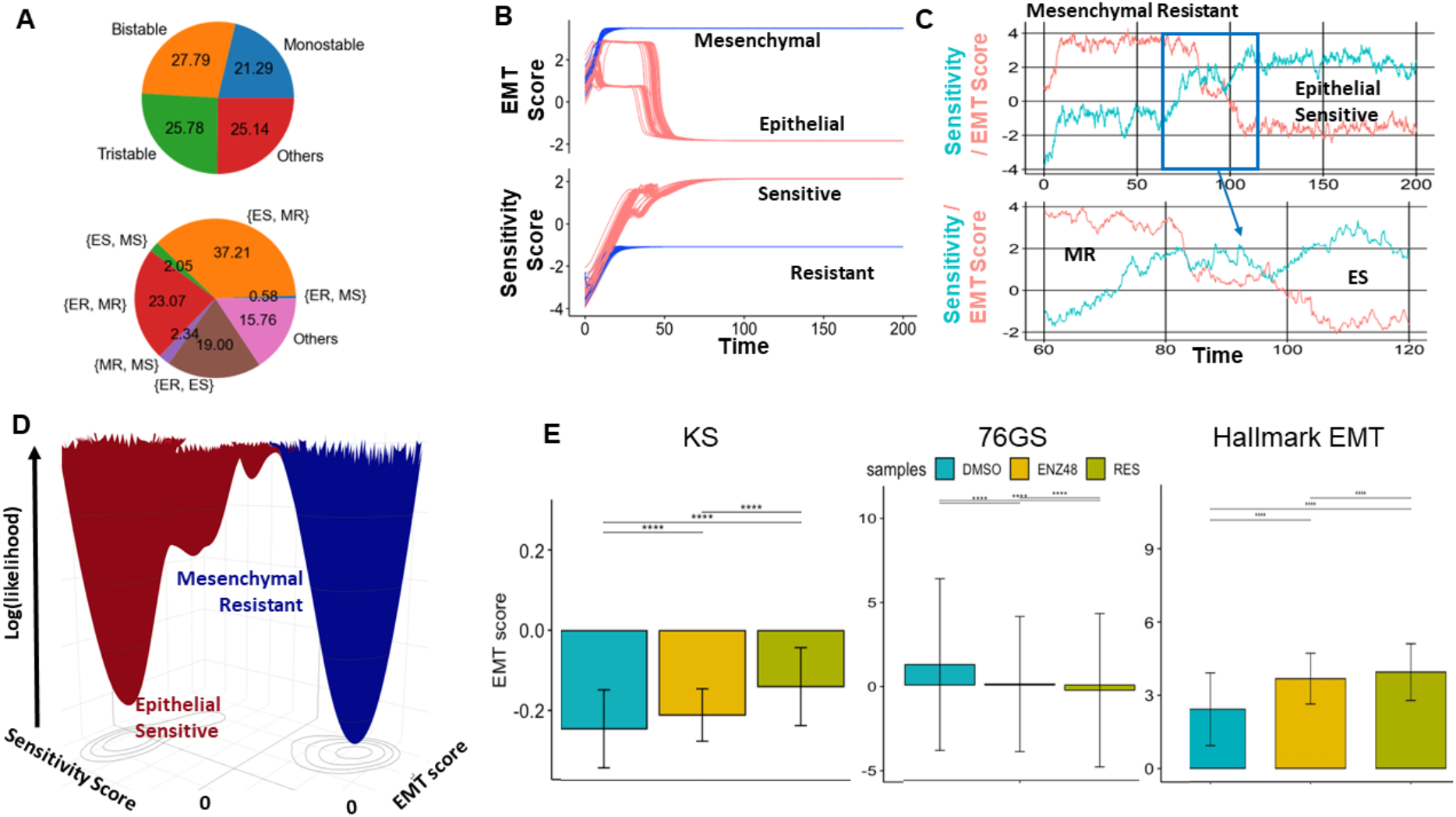
Stochastic cell-state transitions along EMT and AR axes. **A)** (Top) Pie chart showing fraction of RACIPE parameter sets in terms of number of steady-state solutions enabled. (Bottom) Pie chart showing combinations of different phenotypes constituting bistable solutions. **B)** System dynamics for representing two biological EMT phenotypes (E, M and Resistance (R, S)) when starting from multiple different initial conditions. **C)** (Top) Stochastic cell-state transition from MR to ES phenotype under the influence of gene expression noise. (Bottom) A zoomed-in version of highlighted region. **D)** Landscape showing log (likelihood) on the Sensitivity and EM score plane, with the valleys representing the stable states possible in the system - Epithelial Sensitive, Mesenchymal Resistant **E)** EMT scores (KS scores, 76GS scores and ssGSEA scores for the Hallmark EMT gene set) for single-cell RNA-seq dataset (GSE168668) comprising samples (cells) treated with DMSO, treated with Enzalutamide for 48h (ENZ48) and Enzalutamide resistant (RES) cells. ****: p < 0.01 for Students’ t-test.

Next, we examined whether EMT and drug resistance behaviors could drive each other. To do this, we quantified the extent of EMT through three EMT scoring metrics - KS and 76GS scores and ssGSEA scores for Hallmark EMT gene set from single-cell RNA-sequence data for samples treated with dimethyl sulfoxide (DMSO), enzalutamide for 48 hours (ENZ48) and cells resistant to enzalutamide (Res). All three EMT scoring metrics showed very consistent trends in terms of induction of EMT in enzalutamide resistant cells as compared to cells treated with DMSO or acutely treated with enzalutamide for 48 hours (**Fig 3E**). This single-cell transcriptomic data analysis thus revealed a drug-induced transition from epithelial-sensitive to mesenchymal-resistant in prostate cancer cell lines.

### Association of enriched PD-L1 levels with EMT and androgen independence

Another hallmark of tumor aggressiveness is immune evasion, and one key mechanism of immune evasion is through elevated expression of immune checkpoint molecules, such as Programmed Death Ligand I (PD-L1) that can suppress the adaptive immune response. Interestingly, EMT has been reported to be associated with higher levels of PD-L1 and other immune checkpoints (20,61). Therefore, we first examined TCGA data and found a significant positive association between expression levels of PD-L1 and SNA11 **(Fig 4A)**. Using a Snail-inducible human LNCaP95 prostate cancer cell line (62,63), activation of Snail induces upregulation of PD-L1 mRNA **(Fig 4B)** and protein **(Fig 4C)**. This implies that PD-L1 is regulated by the EMT mediator, SNAIL.

**Figure 4:**
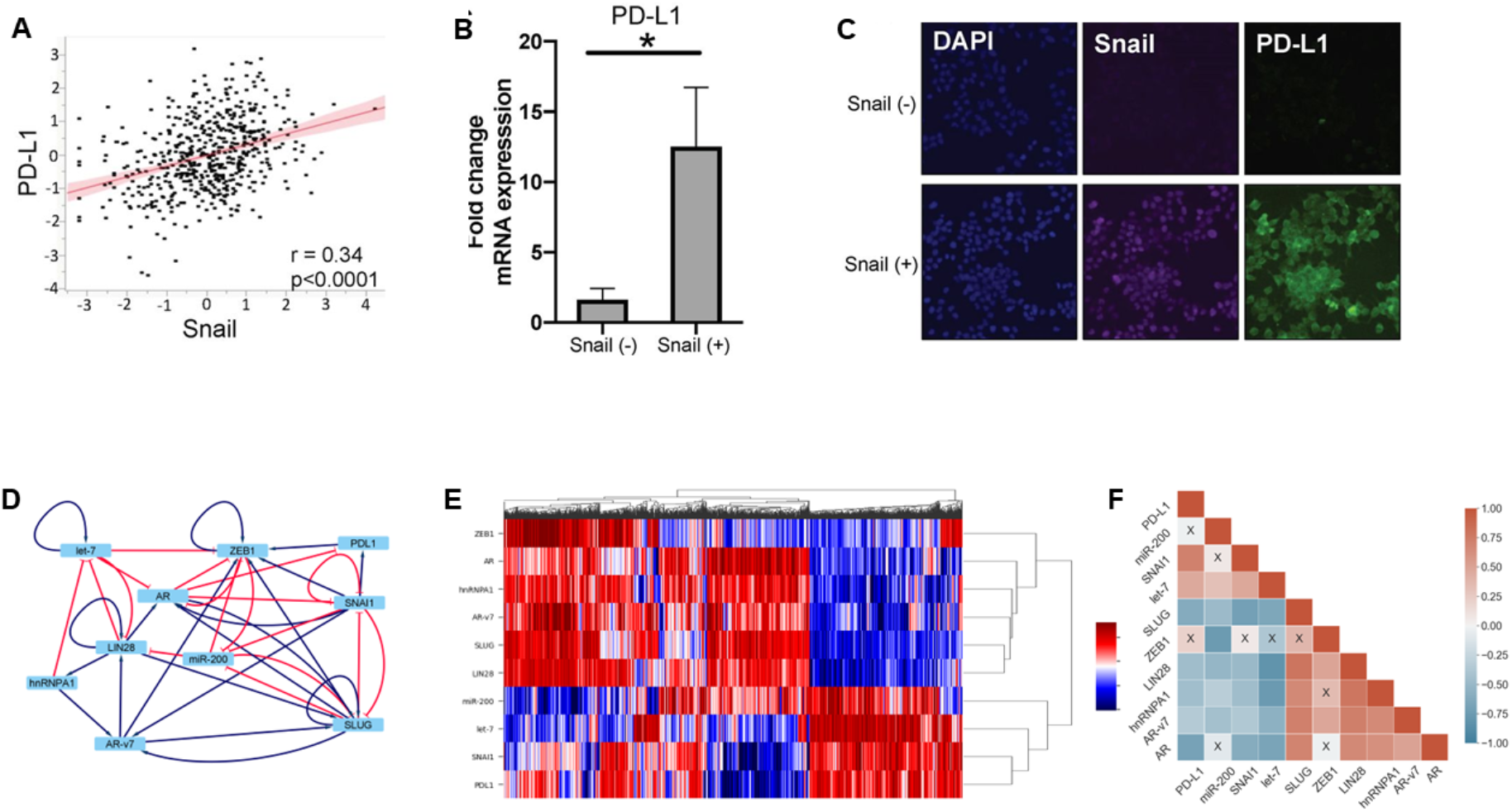
Association of PD-L1 with EMT/AR crosstalk. **A)** RNA-Seq data from The Cancer Genome Atlas shows a significant positive correlation between SNAIL and PD-L1 expression in prostate cancer samples **B)** mRNA expression of PD-L1 with and without SNAIL in the LNCaP95-Snail inducible cell line **C)** Protein expression of SNAIL and PD-L1 in LNCaP95 with and without SNAIL activation **D)** Gene regulatory network showing the androgen receptor axis and EMT related genes, along with an added PD-L1 node **E)** Heatmap of stable steady-state solutions for the network shown in A, obtained via RACIPE **F)** Correlation matrix for the genes in the network; red indicates positive correlation and blue indicates negative correlation; x indicates correlations with p>0.05 and the colour of each cell indicates the strength of the correlation.

PD-L1 is a well-known driver of immune evasion in several cancer types (64). Given the observed association of PD-L1 with the EMT axis and with enzalutamide-resistant prostate cancer cell lines, we expanded our regulatory network to include PD-L1 as a node **(Fig 4D)** and simulated the emergent dynamics of this expanded network. Steady state values obtained from RACIPE simulations for this expanded network were hierarchically clustered in the heatmap **(Fig 4E)**, which showed PD-L1 expression to be closely related to that of SNAI1. Furthermore, high expression of AR coincides with low expression of PD-L1 and *vice versa*, as further corroborated by the pairwise correlation heatmap **(Fig 4F)** obtained from the cohort of RACIPE solutions for this network.

To further investigate these relationships, we probed the correlations between PD-L1 signatures, SNAI1 and AR expression levels in several transcriptomic datasets – TCGA, CCLE, GSE54460, and GSE74685 **(Fig 5A, i; 5B, i))**. These datasets revealed that PD-L1 expression levels significantly correlated positively with SNAI1, but negatively with AR levels. The expression values generated by our network simulations also recapitulated the correlation trends **(Fig 5A,ii; 5B,ii))**.

**Figure 5:**
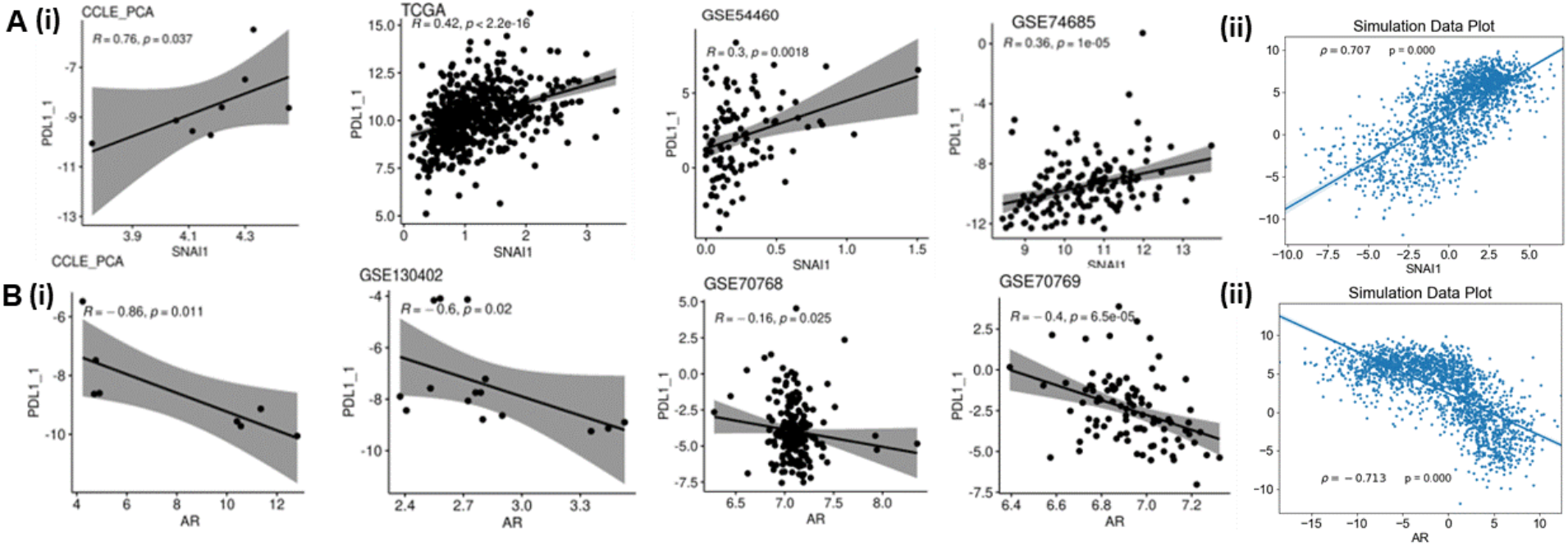
Association of PD-L1 signature with AR in simulation and transcriptomic data. **A) (i)** Scatter plots indicating the Pearson’s correlation between PD-L1 signature (calculated based on ssGSEA scores of 15 genes defined in (66)) and SNAI1 levels in various clinical data sets and **(ii)** Scatter plot indicating the Pearson’s correlation between PD-L1 and SNAI1 levels at steady state produced by simulations **B.(i)** Scatter plots indicating the Pearson’s correlation between PD-L1 and AR levels in various clinical data sets and **(ii)** Scatter plot indicating the Pearson’s correlation between PD-L1 and AR levels at steady state produced by simulations.

To identify whether the above-mentioned trends are evident in additional transcriptomic datasets, we undertook a meta-analysis where we analyzed 70 independent transcriptomic datasets in prostate cancer (**Table S4**). We calculated the ssGSEA scores for epithelial and mesenchymal gene lists, as well as those for Hallmark EMT signature. We observed that across 24 datasets which showed a statistically significant correlation between the epithelial and mesenchymal scores, 22 (~ 92%) of them showed a negative trend, indicating that these two programs are strongly antagonistic to one another (**Fig 6A**, top). The Hallmark EMT scores also correlated strongly positively with Mes scores (**Fig 6A**, bottom). Consistently, we observed a dominance of positive correlation of PD-L1 expression levels as well as Androgen Independence geneset activity with a more mesenchymal phenotype (**Fig 6B–C**). Overall, this meta-analysis (65) confirms the association of EMT and AR-dependent scenario of androgen independence with PD-L1 enrichment in prostate cancer.

**Figure 6:**
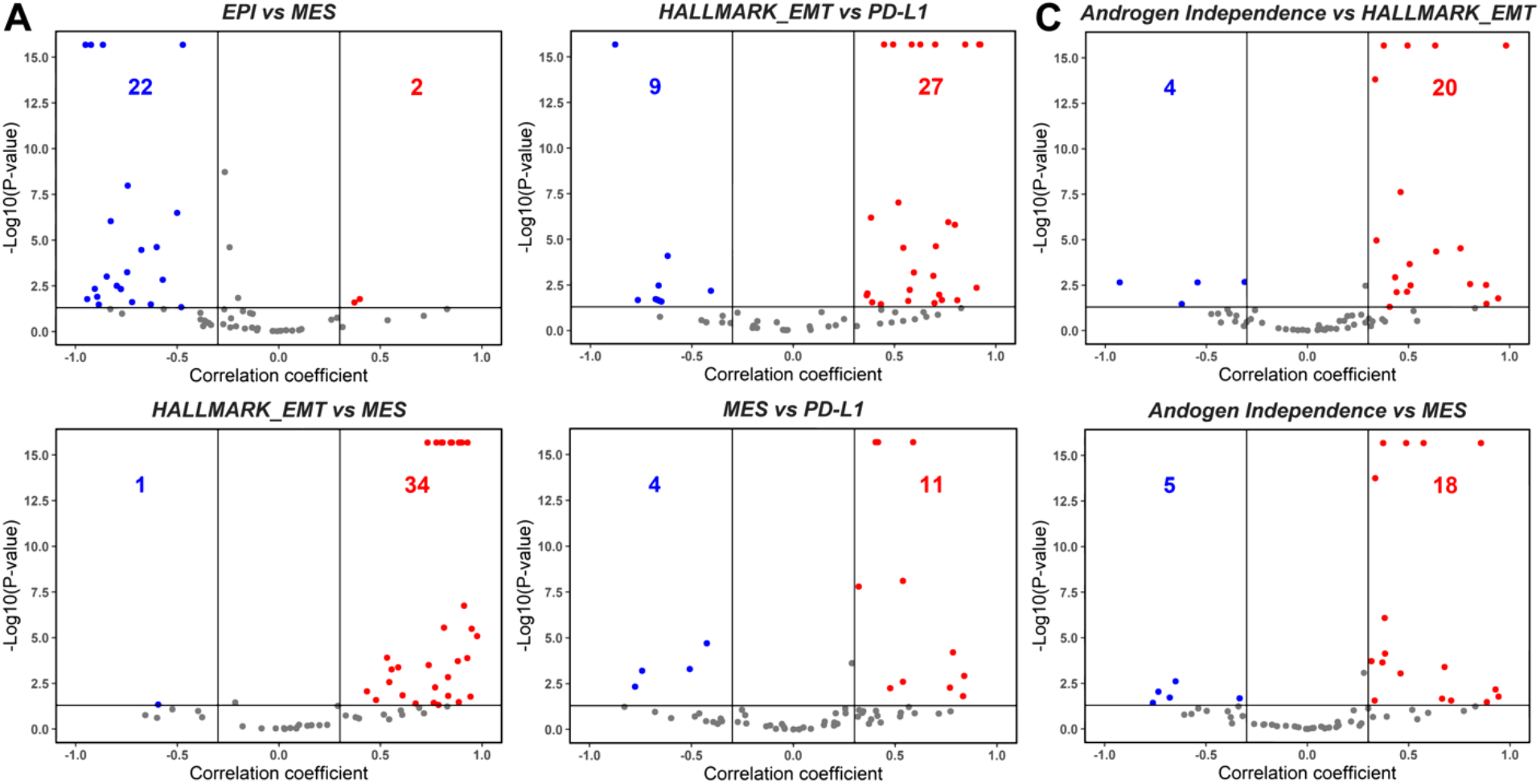
Meta-analysis showing association between EMT, androgen independence and PD-L1 in bulk transcriptomic data. **A)** Volcano plots showing spearman correlation coefficients (x-axis) and -log10(p-value) (y-axis) for Epi vs. Mes (top) and Hallmark EMT vs. Mes (bottom). Significant correlations (R > ± 0.3 and p < 0.05) are shown as red (positive correlation) and blue (negative correlation) datapoints. Each dot represents a unique data set. Numbers on each side of the volcano plots indicate the number of data sets that are positively correlated (red) or negatively correlated (blue). Same as A) but for **B)** Hallmark EMT vs. PD-L1 (top), Mes vs. PD-L1 (bottom), and **C)** Hallmark EMT vs. Androgen Independence (top), Mes vs. Androgen independence (bottom).

## Discussion

EMT-like lineage plasticity is associated with multiple key features of aggressive disease across solid tumors, including metastasis, therapy resistance, and immune evasion. In the current work, we focused on the interconnections of EMT and AR gene regulatory networks in the context of hormone therapy resistance and immune evasion. EMT-like lineage plasticity is associated with multiple key features of aggressive disease across solid tumors, including metastasis, therapy resistance, and immune evasion. In the current work, we focused on the interconnections of EMT and AR gene regulatory networks in the context of hormone therapy resistance and immune evasion.

Here, we adopt a computational systems biology approach to elucidate the mechanisms underlying the development of anti-androgen resistance, by unraveling the emergent dynamics of a minimal gene regulatory network incorporating various experimentally reported interactions among EMT players (Snail, SLUG, miR-200, ZEB1) and signaling by androgen receptor and its variants (AR, AR-v7). The dynamics of this coupled network reveals the (co-) existence of four phenotypes: epithelial sensitive (ES), epithelial resistant (ER), mesenchymal sensitive (MS) and mesenchymal resistant (MR), with ES and MR being the dominant ones. Thus, our model could recapitulate how the transcriptional dynamics of an underlying regulatory network can give rise to phenotypic heterogeneity in terms of both epithelial-mesenchymal status and corresponding anti-androgen resistance traits. Through a careful analysis of underlying network topology, we found that the regulators of EMT and castration resistance organized themselves into “teams” of players which act antagonistic to each other. Such organization of regulatory genes into teams was lost when the edges in this network were shuffled to generate many non-biological randomized hypothetical networks. This observation suggests the evolution of network to this specific topology possibly to reinforce the coupling between EMT and anti-androgen resistance traits. Furthermore, we showed that the ES and MR phenotypes can switch among each other due to noise in gene expression, showcasing phenotypic plasticity features of this network.

These simulation results were endorsed by analysis of multiple transcriptomic datasets both at the bulk and single-cell levels, through demonstrating: a) more mesenchymal cell line or primary tumor (quantified by high KS scores, low 76GS scores, high levels of Zeb1, high ssGSEA scores for Hallmark EMT pathway) associated with an enrichment of the androgen independence geneset in prostate cancer samples, b) enzalutamide-resistant cells had a more mesenchymal nature, and c) PD-L1 expression correlated positively with SNAIL expression but negatively with AR signaling, thus highlighting how EMT/AR crosstalk can impact other axes of plasticity, such as upregulation of immune checkpoints and subsequent immune evasion with potentially important therapeutic implications.

EMT has been previously reported in castration resistant prostate cancer bone metastases (17). Moreover, androgen deprivation has been shown to drive EMT in prostate cancer cell lines and patient derived xenografts (PDXs) (32). Further, over-expression of AR-v7 has been seen to promote induction of mesenchymal and stemness markers (54), thereby strengthening our observations of coupled EMT and anti-androgen resistance axes. Besides the ES and MR phenotypes, our model also predicted two other phenotypes, although at a lower frequency - mesenchymal-sensitive (MS) and epithelial-resistant (ER). Future experiments should investigate the role of these phenotypes and identifying whether these phenotypes are seen especially during MET, given the asymmetric or hysteretic dynamics of EMT/MET. Another model prediction that needs detailed experimental validation is stochastic cell-state transition among cells with varying androgen-dependence.

Our mechanism-based model simulations provide a template of how the emergence of phenotypic heterogeneity can be a function of feedback loops formed by transcriptional crosstalk. However, there are also several limitations to this study. For one, other regulatory layers that act on varying timescales – epigenetic(68), translational (69), genetic (70,71) and metabolic – may need to be incorporated into this framework to better understand their impact on phenotypic plasticity and gene expression heterogeneity and subsequent therapy resistance. Including these additional players and underlying genetic and phenotypic lineages (e.g., TP53/RB1 status, relative AR-dependency, neuroendocrine-like or “double negative” phenotypes) can facilitate more trajectories of anti-androgen resistance, such as lineage plasticity to a neuroendocrine state. Our modeling framework can be expanded to include other reported mechanisms of castration resistance, such as those involving PAGE4 (72) or stemness factors like SOX2 (73). While modeling approaches can always be made more complex, our framework serves as a unique platform to understand the interconnections between multiple gene regulatory axes. This platform can be applied to test combinatorial therapies that can counteract the impact of phenotypic plasticity (similar to efforts for ER+ breast cancer (74)) and/or strategies such as bipolar androgen therapy (75).

## Materials and Methods

### Cytoscape

Cytoscape was used to generate Fig. 1A(i) and 4A(i). A table containing source node, target node and the type of interaction (activation or inhibition) was provided as input to generate these networks.

### RACIPE (<u>R</u>andom <u>C</u>ircuit <u>P</u>erturbation)

Random Circuit Perturbation (RACIPE)(76) is a computational tool that takes the network topology (nodes and interactions) as input and generates an ensemble of kinetic models with parameters and initial conditions sampled within biologically relevant ranges. Each such model consists of coupled ordinary differential equations (ODEs) for each node and is represented as the basal production rate multiplied with shifted Hill functions for each interaction of which the gene is a target, along with a degradation term. These ODEs were solved using Euler’s method for multiple sets of initial conditions to give steady states. The ensemble of steady states that are obtained from this represent the variability in expression by different cells in a population. For our simulations, we used 10,000 parameter sets (10,000 models), where each model was solved for 100 initial conditions.

### Z-normalization

The RACIPE stable state data generated is log2 normalized. We performed z-score normalisation on the data using the expression given below:

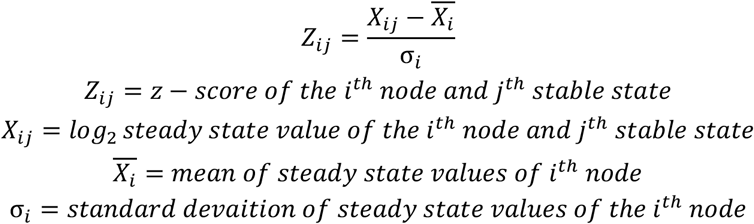

### Kernel Density Estimate

This method is used to estimate the probability density function by smoothening the data of a random variable, which in our case is the steady state levels for each node. We used the function *kdeplot* under the library *seaborn* in Python 3.9.5 to generate the plots. Histograms were generated to show the discrete probability density function, with number of bins fixed at 24. Bimodality coefficients were calculated for each distribution using Sarle’s bimodality index (described later).

### Correlation Plots

To calculate correlations, we used Spearman correlation coefficients and the corresponding p-values to assess the statistical significance of these correlations. For the correlation between Snail and PD-L1 from The Cancer Genome Atlas prostate cancer (PRAD) data set, the correlation between Snail expression and PD-L1 expression from prostate cancer patients was analyzed using the Kruskal Wallis test.

### Finding Boundary Conditions for determination of Phenotypes

To classify the steady state solutions into biologically meaningful cell states, we calculated the Epithelial-Mesenchymal (EM) scores and AR Resistance scores for each sample in the simulated dataset, which help us determine quantitatively the phenotype a certain steady state corresponds to. The difference between the Z-normalized steady state values of ZEB1 and miR200 was used to calculate the EM score while the sum of the Z-normalized steady state values of AR and AR-v7 were used to define the Resistance score. Both EM and Resistance scores were z-normalized across all simulated samples. On plotting the kernel density estimates of these scores, we found EM and resistance to be bimodal **(Fig. C(i), D(i))**. Thus, each score corresponded to two phenotypes – epithelial and mesenchymal for EM score and sensitive and resistant for Resistance score. To define the boundary score between two phenotypes, we calculated the central minima between the two peaks of the kernel density estimate for the scores determined from the biological network.

### Uniform Manifold Approximation and Projection (UMAP)

UMAP is a dimensionality reduction method which is based on topological analysis of the data. The n-dimensional data is reduced to 2 dimensional UMAP coordinates. For Fig. 1C(ii) and 1D(ii) each point is coloured based on Resistance scores and EM scores, respectively. We used the *umap-learn* library in Python to generate this plot. The n_neighbors parameter which determines the number of nearest neighbours to consider while the clustering was set to 100. The min_dist parameter which determines the region to look at for clustering was set to 0.8.

### Principal Component Analysis

PCA is a dimensionality reduction tool which gives us an idea of the multidimensional network. To analyse the degree to which dimensionality of the system could be reduced, we looked at the Scree plot which showed that PCA1 and PCA2 could explain the major features of the network (**Fig 2C (i)**).

### Generation of Random Networks

To generate unique randomized networks for the ‘wild type’ network, the directed edge topology of the biological network was kept intact. Hence, the source and target of each edge remained the same. However, each edge was assigned as activating or inhibiting randomly such that the overall number of activating edges and inhibiting edges remained the same as that in the ‘wild type’ network. We generated an ensemble of 100 of such randomized networks for the analysis (Supplementary Figure 4).

### Bimodality Coefficient (BC)

BC assumes that the distribution is unimodal as its null hypothesis. For values more than 0.555, we interpret that the null hypothesis has failed, and the distribution is not unimodal. The bimodality coefficient was calculated using the formula given below:

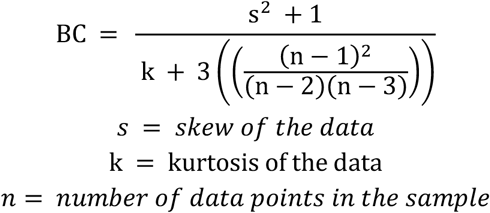

### Hartigan’s dip test (HDT)

The HDT assumes that the unimodal hypothesis as the null hypothesis(77). Tests with a p-value <0.05 confirmed that the distribution is not unimodal. The *diptest* package in R was used to test for bimodality.

### Influence Matrix and Team Strength

The influence matrix was calculated as defined previously in our work (39). It quantifies the effect each node has on another node solely based on the topology of the network. For a given pair of nodes, it defines a path as the collection of serially connected edges that start from one of the nodes and ends at the others and defines the pathlength as the number of edges in the path.

Team strength is a metric which quantifies the strength of the two teams seen in the influence matrix, in other words, how strongly the members within a team support each other and those across teams inhibit each other. It is defined on a scale of [0, 1]; the higher the value, the more well-defined the two teams. This is calculated by the formula defined in (78).

### Datasets analyses

For bioinformatic analysis publicly available datasets CCLE(79), TCGA (https://www.cancer.gov/tcga.), GSE54460(80), GSE74685(81), GSE707768, GSE70769(82), GSE77959(52), GSE80042(45), GSE67681(53), GSE22010(83), GSE 168668 and GSE130402 were used. The results published here are in part based upon data generated by The Cancer Genome Atlas (TCGA) Research Network: http://cancergenome.nih.gov.

### Gene signatures

To calculate gene signatures for each sample, we used single sample GSEA (ssGSEA) (84) scores for a given gene set. AR signature scores were calculated based on the WANG_PROSTATE_CANCER_ANDROGEN_INDEPENDENT geneset(29) and EMT signature scores were calculated using HALLMARK_EPITHELIAL_MESENCHYMAL_TRANSITION geneset available on MsigDB. EMT scores KS and 76GS were calculated based on previous reports (85,86). PD-L1 signature was calculated based on ssGSEA scores of 15 genes defined previously in literature (66).

### Stochastic simulations

We simulated network in described in Figure 1 A (i) using Euler–Maruyama method for some representative parameter sets (which were taken from RACIPE) that corresponded to coexistence of two states high EM score-low sensitivity score (Mesenchymal Resistant) and low EM score-high sensitivity score (Epithelial sensitive). The corresponding equations take the following form:

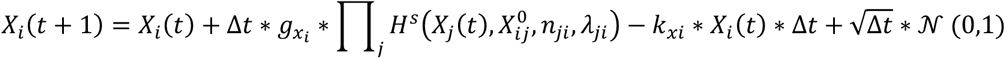

The equation used for stochastic simulations here are similar to ODEs used by RACIPE but with noise terms: 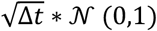,where △t is the time step and 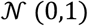 is a normal random variable with mean 0 and standard deviation 1. For the parameter set, we simulated the network for 100 different initial conditions sampled uniformly from the range 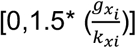. We then normalized the trajectories using the mean and standard deviation of each node expression obtained from RACIPE and converted the trajectories to EM scores and PD-L1 levels in order to classify them into the observed phenotypes. Using these trajectories, we constructed obtained a probability density (P) of the EM-Sensitivity score pairs and constructed a log (likelihood) landscape (87).

### Cell culture

LNCaP95-Snail inducible cells were cultured in RPMI containing 10% charcoal-stripped fetal bovine serum and 1% penicillin/streptomycin and maintained at 37°C in a humidified incubator with 5% CO_2_. Induction of Snail nuclear translocation was mediated by the addition of 4-hydroxy-tamoxifen (40HT) at a concentration of 20 nM. Ethanol (EtOH) was used as a vehicle control. All cells were authenticated by the Duke DNA Analysis Facility using analysis of short tandem repeats and were verified to be mycoplasma-free.

### Real-Time quantitative RT-PCR

For qPCR, total RNA was reverse transcribed using the High-Capacity cDNA Reverse Transcription Kit (Life Technologies). Aliquots of 5-fold diluted reverse transcription reactions were subjected to quantitative (q)PCR with KAPA SYBR FAST master mix using the Vii7 real time-PCR detection system (Applied Biosystems). GAPDH mRNA levels were measured for normalization, and the data are presented as “Relative Expression”.

### Immunofluorescence staining

For cells expressing inducible Snail, cells were pretreated with ethanol (EtOH) or 4OHT. For immunofluorescence (IF) staining, cells were fixed in 4% PFA, permeabilized with 0.2% Triton X-100, and stained with Hoechst. Cells were blocked with 5% bovine serum albumin (BSA, Sigma) prior to incubation with primary antibodies. Cells were incubated in Alexa Fluor secondary antibodies (Life Technologies) and then imaged on an inverted Olympus IX 73 epifluorescence microscope.

## Data Availability

All codes are available publicly at https://github.com/csbBSSE/AR_Prostate_Cancer.

## Author contributions

RJ, AN, MP, KW, DS and MS performed research and contributed to data generation. JAS and MKJ conceptualized and supervised research. All authors contributed to data analysis and writing and review of the manuscript.

## Supplementary Figures

**Fig S1:.**
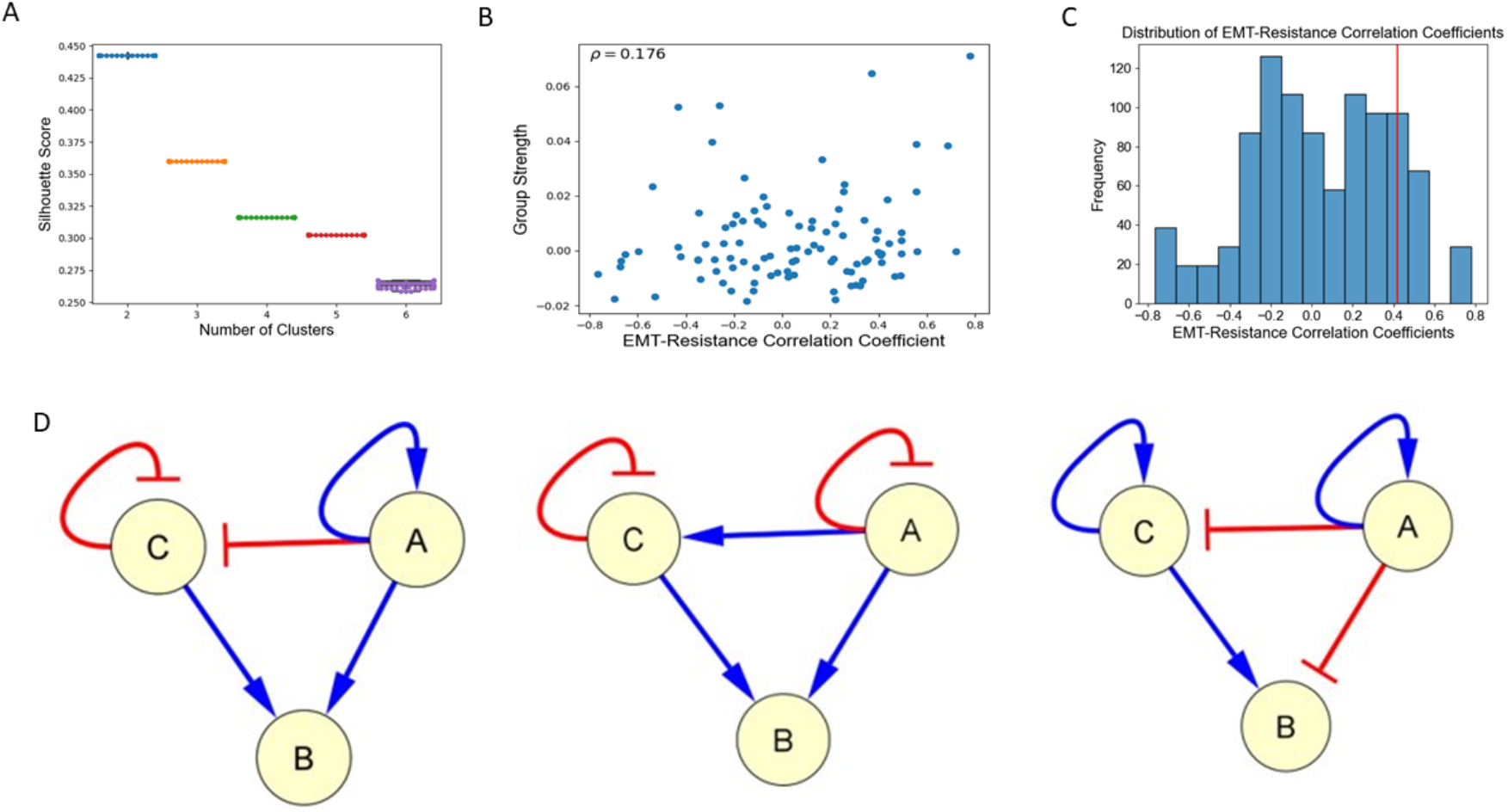
**A.** Silhouette score boxplots made for 100 iterations of k-means clustering for each integer number of clusters (K=2-6). **B.** Scatter plot of EM-Resistance Spearman’s correlation coefficients for an ensemble of 100 randomized networks. **C.** Histogram of the EM-Resistance Spearman’s correlation coefficients for an ensemble of 100 randomized networks. **D.** Generating a random network by keeping the number of inhibitions and activations fixed, the red arrow indicates inhibition with arrowhead pointing to target node; blue arrow indicates the activation with arrowhead pointing towards the activation.

**Fig S2.**
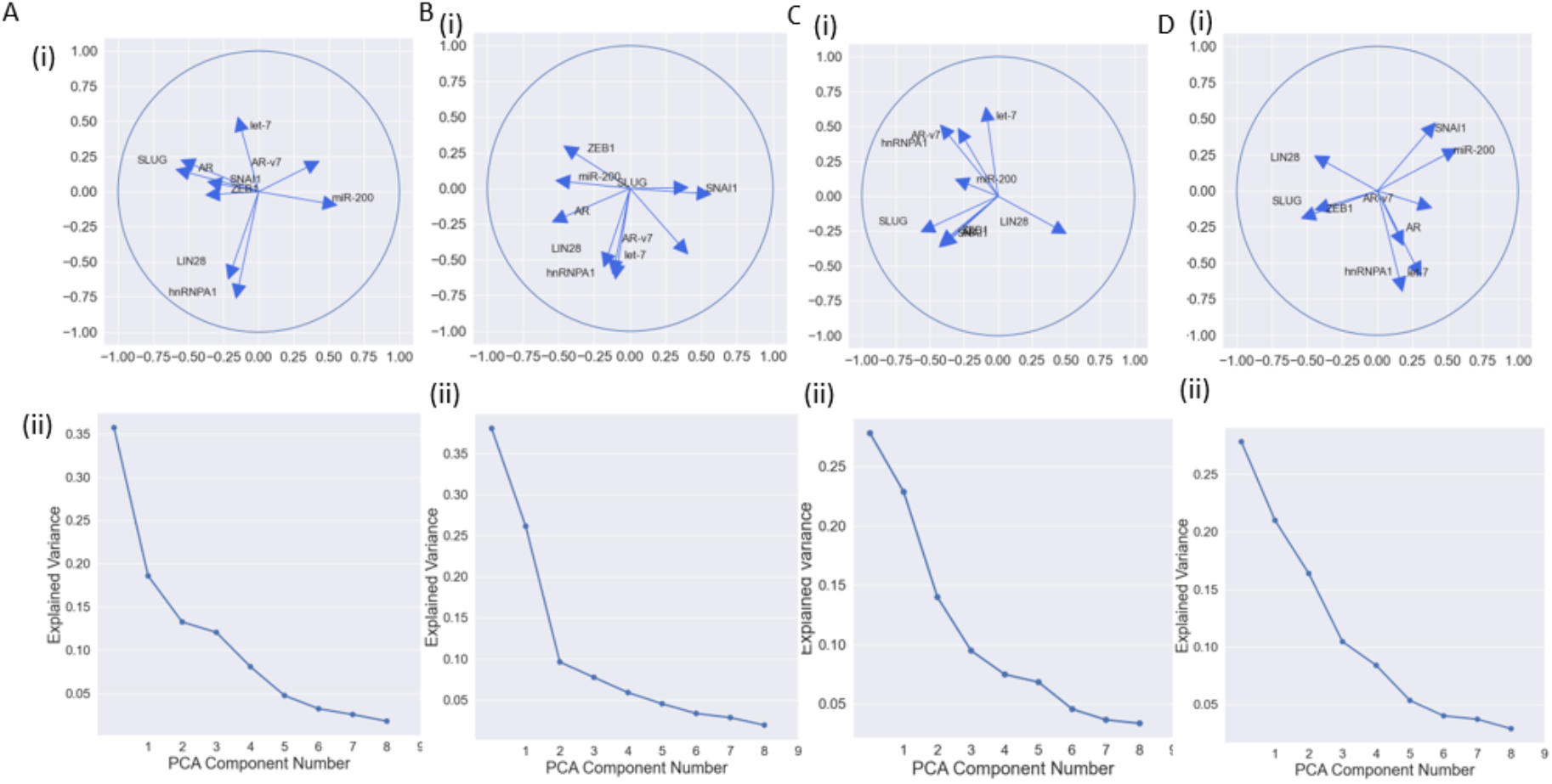
A-D) (i) Scree plot showing PCA components and their explained variance ratio (above) and (ii) PCA Correlation Circle of the RACIPE solutions of a random biological network.

