## Supplementary material for "Emergent dynamics of underlying regulatory network links EMT and androgen receptor-dependent resistance in prostate cancer": Tables S1-S3

| **Source** | **Target** | **Type of regulation** | **Level of Regulation** | **Type of interaction** | **Reference** |
| --- | --- | --- | --- | --- | --- |
| ZEB1 | miR-200 | Inhibition | E-box binding site on miR-200 | Direct | (1,2) |
| miR-200 | ZEB1 | Inhibition | Transcriptional due to presence of binding site | Direct | (3) |
| SNAI1 | SNAI1 | Inhibition | Transcriptional level auto-regulation | Direct | (4) |
| SNAI1 | ZEB1 | Activation | Transcriptional level regulation | Direct | (5) |
| AR | SLUG | Activation | At the mRNA levels |  | (6,7) |
| SLUG | AR | Activation | DBD region of AR is where SLUG binds | Direct | (6,7) |
| let-7 | LIN28 | Inhibition | miRNA-mediated translational repression | Direct | (8,9) |
| LIN28 | let-7 | Inhibition | Both Drosha and Dicer levels of let 7 biogenesis, inhibition of microRNA processing | Direct | (8–10) |
| let-7 | let-7 | Activation | Auto-catalysis of miRNA processing | Direct | (11) |
| miR-200 | LIN28 | Inhibition | Transcriptional regulation, the members of miRNA family have binding sites on Lin28B gene | Direct | (12) |
| let-7 | AR | Inhibition | Upregulation at mRNA levels |  | (13,14) |
| AR | SNAI1 | Inhibition | Transcriptional mediated by DHT | Indirect | (15) |
| SLUG | SLUG | Activation | Transcriptional Activation | Direct | (16) |
| hnRNPA1 | let-7 | Inhibition | Drosha level of maturation of let-7 | Direct | (17) |
| hnRNPA1 | AR-v7 | Activation | pre-mRNA splicing - promotes splicing at AR V7 splice sites | Direct | (18) |
| SNAI1 | AR | Activation | Upregulation data with a binding site present, binding data not shown |  | (19) |
| SLUG | ZEB1 | Activation | Transcriptional at E box binding site (In Melanoma Cell line) | Direct | (20) |
| ZEB1 | ZEB1 | Activation | Via SMAD | Indirect | (21–23) |
| ZEB1 | AR | Inhibition | Transcriptional suppressor of AR ENCODE identified 3 Zeb1-binding sites in AR genomic sequences | Direct | (24) |
| AR | ZEB1 | Inhibition | Knockdown based | Indirect | (24) |
| LIN28 | AR | Activation | Upregulation and shLin data support activation |  | (25) |
| SLUG | miR-200 | Inhibition | Indirect via TGF-B but has binding site on miR200 | Indirect | (26) |
| SNAI1 | miR-200 | Inhibition | Transcriptional Repressor |  | (27) |
| miR-200 | SLUG | Inhibition | Binding site at 3'UTR region of SLUG gene | Direct | (26) |
| LIN28 | LIN28 | Activation | Translational activation | Direct | (28,29) |
| SNAI1 | AR-v7 | Activation | Upregulation data with a binding site present, binding data not shown |  | (19) |
| SLUG | AR-v7 | Activation | ARE activity increased |  | (6) |
| AR-v7 | ZEB1 | Activation | Upregulation data |  | (30) |
| let-7 | ZEB1 | Inhibition | HMGA2 mediated | Indirect | (31) |
| LIN28 | hnRNPA1 | Activation | Upregulation, not known at what levels |  | (25,32) |
| LIN28 | SLUG | Activation | shLin decreased SLUG levels thereby implying activation of SLUG |  | (9) |
| AR-v7 | SLUG | Activation | Regulation at mRNA levels |  | (6) |
| SLUG | SNAI1 | Inhibition |  |  | (33) |
| SNAI1 | SLUG | Inhibition |  |  | (33) |
| Ar-v7 | LIN28 | Activation | Upregulation data supports the activation |  | (30) |

**Table S1**: The table consists of Source node in Column one and Target node in Column 2, with type of interaction mentioned in column 3 and column 4 and 5 consist of short description of type of interaction and its origin, column 5 has the references.

| **Genes** | **Run A** | **Run B** | **Run C** | **Mean Value** |
| --- | --- | --- | --- | --- |
| ZEB1 | 0.000 | 0.000 | 0.000 | 0.000 |
| miR-200 | 0.593 | 0.675 | 0.983 | 0.751 |
| SNAI1 | 0.995 | 0.973 | 0.998 | 0.989 |
| AR | 0.975 | 0.997 | 0.993 | 0.989 |
| SLUG | 0.000 | 0.000 | 0.000 | 0.000 |
| let-7 | 0.000 | 0.000 | 0.000 | 0.000 |
| LIN28 | 0.000 | 0.000 | 0.000 | 0.000 |
| hnRNPA1 | 0.000 | 0.000 | 0.000 | 0.000 |
| AR-v7 | 0.988 | 0.994 | 0.810 | 0.931 |

**Table S2**: The Hartington’s Diptest Table consists of HDT value for 3 different Runs with their mean values in the Last column.

| **Genes** | **Run A** | **Run B** | **Run C** | **Mean Value** |
| --- | --- | --- | --- | --- |
| ZEB1 | 0.58 | 0.576 | 0.578 | 0.581 |
| miR-200 | 0.434 | 0.433 | 0.436 | 0.435 |
| SNAI1 | 0.439 | 0.427 | 0.423 | 0.430 |
| AR | 0.440 | 0.439 | 0.442 | 0.441 |
| SLUG | 0.644 | 0.638 | 0.644 | 0.642 |
| let-7 | 0.636 | 0.625 | 0.627 | 0.630 |
| LIN28 | 0.676 | 0.665 | 0.667 | 0.669 |
| hnRNPA1 | 0.580 | 0.576 | 0.579 | 0.578 |
| AR-v7 | 0.426 | 0.428 | 0.422 | 0.425 |

**Table S3:** consists of Bimodality Coefficient for different Runs and their Mean values in the last column. BC<0.555 is used to reject null hypothesis stating the component is not bimodal.
